# Computational Exploration of Protein Structure Dynamics and RNA structural Consequences of *PKD1* Missense Variants: Implications in ADPKD Pathogenesis

**DOI:** 10.1101/2024.03.21.586139

**Authors:** Chandra Devi, Prashant Ranjan, Sonam Raj, Parimal Das

**Affiliations:** Centre for Genetic Disorders, Institute of Science, Banaras Hindu University, Varanasi, U.P., India-221005; National Cancer Institute, National Institutes of Health, Bethesda, Maryland-20892

**Author notes:** Correspondence Centre for Genetic Disorders, Institute of Science, Banaras Hindu University, Varanasi, U.P., India-221005.

**Keywords:** ADPKD, MD Simulation, PKD1, missense, Protein dynamics, RNA structure

## Abstract

Autosomal dominant polycystic kidney disease (ADPKD), a genetic disorder characterized by the formation of fluid-filled cysts within the kidneys, leading to progressive renal dysfunction, is primarily caused by mutations in *PKD1*, a gene encoding for the protein polycystin-1 (PC1). Understanding the structural consequences of *PKD1* variants is crucial for elucidating disease mechanisms and developing targeted therapies. In this study, we analyzed the effects of nine missense *PKD1* variants, including c.6928G>A p.G2310R, c.8809G>A p.E2937K, c.2899T>C p.W967R, c.6284A>G p.D2095G, c.6644G>A p.R2215Q, c.7810G>A p.D2604N, c.11249G>C p.R3750P, c.1001C>T p.T334M, and c.3101A>G p.N1034S on RNA structures, their interactions utilizing computational tools. We also explain the effects of these variants on PC1 protein dynamics, stability, and interactions using molecular dynamics (MD) simulation. These variants are located at crucial domains such as the REJ domain, PKD domains, and cation channel domain, potentially compromising PC1’s function and contributing to ADPKD pathogenesis. Findings reveal substantial deviations in RNA structures and their interactions with other proteins or RNAs and also protein structure and dynamics for variants such as c.8809G>A (p.E2937K), c.11249G>C (p.R3750P), c.3101A>G (p.N1034S), c.6928G>A (p.G2310R), c.6644G>A (p.R2215Q) suggesting their potential implications in disease etiology. The study also suggests that although certain variants may have minimal effects on RNA conformations, their observed alterations in MD simulations indicate potential impact on protein structure dynamics highlighting the importance of evaluating the functional consequences of genetic variants by considering both RNA and protein levels. This study offers valuable perspectives of the utility of studying the structure dynamics through computational tools in prioritizing the variants for their functional implications and understanding the molecular mechanisms underlying ADPKD pathogenesis and developing therapeutic interventions.

**GRAPHICAL ABSTRACT:** 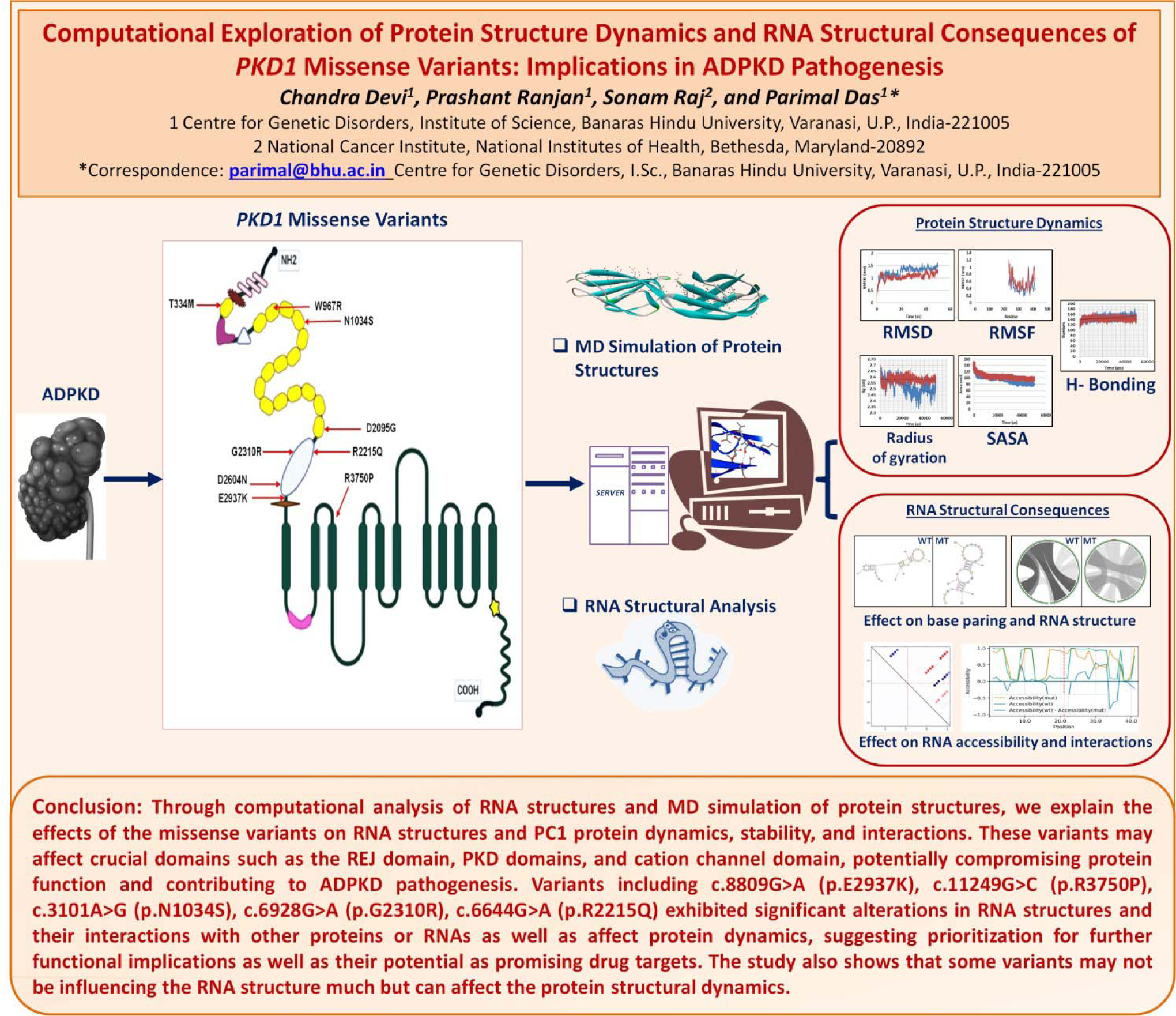

## Introduction

Polycystic kidney disease (PKD) stands as a formidable challenge in the realm of renal disorders, with its hallmark manifestation being the formation of fluid-filled cysts in the kidneys, often leading to progressive renal dysfunction and, eventually, end-stage renal disease (ESRD) in severe cases. Autosomal dominant PKD (ADPKD) is the most common form of PKD, affecting approximately 1 in 1000 individuals worldwide. It is primarily caused by mutations in two genes, *PKD1* and *PKD2*, encoding for the proteins polycystin-1 (PC1) and polycystin-2 (PC2), respectively (Hopp et al., 2020; Yeung et al., 2024). The full function of this protein is not entirely clear, but PC1 is known to be a large transmembrane protein with a complex structure and crucial role in regulating cellular processes such as differentiation, cell proliferation, and apoptosis (Peintner & Borner, 2017). It comprises an N-terminal extracellular domain, several transmembrane domains, and a cytoplasmic C-terminal tail. The leucine-rich repeat (LRR) domains are implicated in signal transduction pathways and mediate PC1’s involvement in cell-cell and cell-matrix interactions. The C-type lectin domain suggests roles in protein-protein interactions and cell adhesion. The sixteen PKD repeats are considered vital for cell-cell interactions and normal renal development. The REJ domain regulates PC1’s ion transport, while the PLAT domain facilitates protein-protein and protein-lipid interactions in signaling pathways. The 11 transmembrane domains likely serve as channels for ion transport, and the C-terminal tail regulates downstream signaling pathways through interactions with G protein subunits, highlighting the important functional roles of PC1 in cellular physiology. PC1 present in primary cilia in the renal tubules are believed to perceive fluid movement within these tubules, contributing to the maintenance of their size and structure. The PC1 and PC2 complex within renal tubules facilitates the typical development and operation of the kidneys (Wang et al., 2019; Weston et al., 2003). Mutations in *PKD1* are responsible for around 78% of ADPKD cases, making it a primary focus for understanding disease pathogenesis (Hopp et al., 2020). Due to the limited understanding of the disease effective cure still remains a challenge. Understanding the structural and functional consequences of specific *PKD1* variants is essential for elucidating genotype-phenotype correlations and guiding personalized treatment approaches.

The experimental determination of its full thermodynamic structural changes at RNA and protein level, and the complete protein’s function remain challenging due to its large complex structure and transmembrane localization. Molecular dynamics (MD) simulation techniques provide valuable insights in studying the dynamic behaviour, structural stability of proteins at the atomic level and interactions within the cellular milieu like how they undergo conformational changes (Hollingsworth & Dror, 2018; Vander Meersche et al., 2024). By computationally modeling the interactions between atoms and molecules over time, MD simulations can help understand the impact of mutations on protein structure, conformational dynamics, and interactions with ligands or other biomolecules making them plausible targets for drug development after understanding the disease mechanisms (Hollingsworth & Dror, 2018; Salo-Ahen et al., 2020). PKD is a complex disease and its complexity can be seen at every scientific level, be it genetic, proteomic or how the disease develops and affects patients. While much research is being focused on understanding the genetics and protein-level consequences of the mutations, understanding their impact on RNA structure and dynamics remains comparatively less explored. There is a need to deepen our understanding at each level of the molecular processes underlying PKD pathogenesis. RNA serves as the intermediary between DNA and protein and plays a crucial role in the regulation of gene expression and hence cellular processes. RNA molecules not only fold into secondary structures but also in three dimensions. The way RNA works depends a lot on its shape, which is influenced by its sequence. The RNA often has to fold many times to get the structure which is a complex process. Usually, RNA starts by folding into simpler shapes that are most energetically favorable, then it forms a paired double helix by folding on itself (Butcher & Pyle, 2011; Draper et al., 2005; Ganser et al., 2019; Holbrook, 2008). Mutations in genes, subsequently within RNA sequences can disrupt its folding and hence structure, affecting its function and potentially contributing to disease pathogenesis (Diederichs et al., 2016; Halvorsen et al., 2010; Hunt et al., 2014; Salari et al., 2013; Sauna & Kimchi-Sarfaty, 2011). Understanding how genetic variants affect RNA could also provide insights into the molecular mechanisms driving PKD progression and potential therapeutic targets. In this study, as we explore into RNA behaviour affected by the missense *PKD1* variants previously identified in ADPKD patients, uncovering another layer of complexity, highlighting RNA’s importance in understanding the disease more thoroughly. We also utilized MD simulation to analyze their structure dynamics and functional effects at protein level. Our results contribute to enhance our understanding of *PKD1*-related disease mechanisms and could potentially guide the development of new therapeutic approaches for managing ADPKD.

## Methodology

### *PKD1* Variants

The wild-type and missense variants of *PKD1* c.6928G>A p.G2310R, c.8809G>A p.E2937K, c.2899T>C p.W967R, c.6284A>G p.D2095G, c.6644G>A p.R2215Q, c.7810G>A p.D2604N, c.11249G>C p.R3750P, c.1001C>T p.T334M, and c.3101A>G p.N1034S, identified in our previous studies using Sanger sequencing and whole exome sequencing were studied for RNA structure using available online tools and protein dynamics using MD simulation. These identified missense variants were individually found in different patients diagnosed with ADPKD (DEVI et al., 2024; Raj et al., 2020).

### RNA Sequence Selection and Extraction

The RNA sequences of interest were derived from the *PKD1* NM_001009944 transcript ID. Short RNA snippets comprising 41 nucleotides were extracted for each wild type and mutant, with the mutation locus precisely positioned at the 21st nucleotide, ensuring 20 bases on both sides of the nucleotide change.

### Prediction of RNA Secondary Structure

The secondary structures of each wild-type and mutant RNA snippets were predicted utilizing the RNAStructure Web Server (version 6.0.1) (Bellaousov et al., 2013). Using thermodynamic principles, this server employs algorithms to predict RNA secondary structures and provides the folding patterns and base pairing interactions within the RNA molecule.

### Effect of Mutation on RNA

Mutational analysis of RNA snippets was done using MutaRNA tool to analyze the structural changes induced by the each missense mutation (Miladi et al., 2020). This analysis involved the intra-molecular base pairing potential, base pairing probabilities of the mutant RNA, and assessment of accessibility (single-strandedness) in comparison to the wild-type counterpart (Bernhart et al., 2011). By combining the remuRNA (Salari et al., 2013) and RNAsnp this tool helps to understand of mutation-induced alterations in RNA structures.

### Mutant Protein Structure Creation and Preprocessing

Mutant structures of PKD1 protein variants were generated utilizing the Swiss Model (Schwede et al., 2003). Given the substantial size of the PKD1 protein, conventional structure modelling and simulation posed significant challenges. Therefore, in lieu of direct structure determination, we employed motif and domain analysis of PKD1 through the motif scan web server (Sigrist et al., 2010). Then we identified location of the variants and subsequently created mutations within the domain structure of PKD1.

Individually, 9 variant structures were generated based on their specific locations within domains and motifs simultaneously the production of 9 distinct wild-type PKD1 structures. To facilitate modeling, different template structures were selected based on the mutation’s location within different domains. The selection of template IDs (T.I.) was as follows: p.E2937K (T.I. G9KGT4.1.A_European domestic ferret), p.G2310R (T.I. H3BTE0.1A_Human), p.W967R (T.I. A0A212CYZ4.1.A_European Red deer), p.D2095G (T.I. Q59EY6.1_Human), p.R2215Q (T.I. H3BTE0.1A_Human), p.D2604N (T.I. H3BTE0.1A_Human), p.R3750P (T.I. 6a70.1b_Human), p.T334M (T.I. A0A212CZX9.1.A_ European Red deer), and p.N1034S (T.I. A0A212CYZ4.1A_European Red deer).

Preprocessing involved the elimination of all non-standard residues, including water molecules, followed by the addition of hydrogen atoms using the Discovery Studio program (Systèmes, 2016). Monomeric structures of PC1 were then isolated for further analysis, with additional non-standard residues removed. Energy minimization of mutant structures was subsequently conducted utilizing Modrefiner (Xu & Zhang, 2011).

### Molecular Simulation Dynamics

MD simulations of wild-type and mutant PC1 regions were conducted using GROMACS (GROMACS96 54a7 force field) (Ranjan & Das, 2023; Van Der Spoel et al., 2005). The systems were solvated with spc water models in a triclinic box. The systems were neutralized by adding Na+ and Cl-ions, and the salt concentration was maintained at 0.15M. The protein was kept at least 1.0 nm from the box edges. Energy minimization was performed using the steepest descent algorithm, followed by equilibration at 300K temperature and 1 atm pressure using the NVT (constant number of particles, volume, and temperature) and NPT (constant number of particles, pressure, and temperature) ensembles. MD simulations were extended to 50ns time frame, and analysis of Root Mean Square Deviation (RMSD), Root Mean Square Fluctuation (RMSF), radius of gyration (Rg), H-bonding, solvent accessible surface area (SASA), was conducted using GROMACS tools. Visualization of MD trajectory data was performed using the XMGRACE application and MS excel (Cowan & Grosdidier, 2000).

## Results

### RNA Secondary Structure

RNA secondary structures predicted using the RNAStructure web server to study the secondary structural changes are depicted in figure 2. The analysis of secondary structure changes due to *PKD1* missense variants revealed prominent alterations in case of variants c.6928G>A (p.G2310R), c.8809G>A (p.E2937K), c.6644G>A (p.R2215Q), c.11249G>C (p.R3750P), and c.3101A>G (p.N1034S). In these variants, significant deviations from the wildtype secondary structure were observed, indicating impact on RNA folding and stability. Conversely, variants c.2899T>C (p.W967R), c.6284A>G (p.D2095G), c.7810G>A (p.D2604N), and c.1001C>T (p.T334M), exhibited less pronounced changes in secondary structure, suggesting milder effects on RNA conformation.

**Figure 1:**
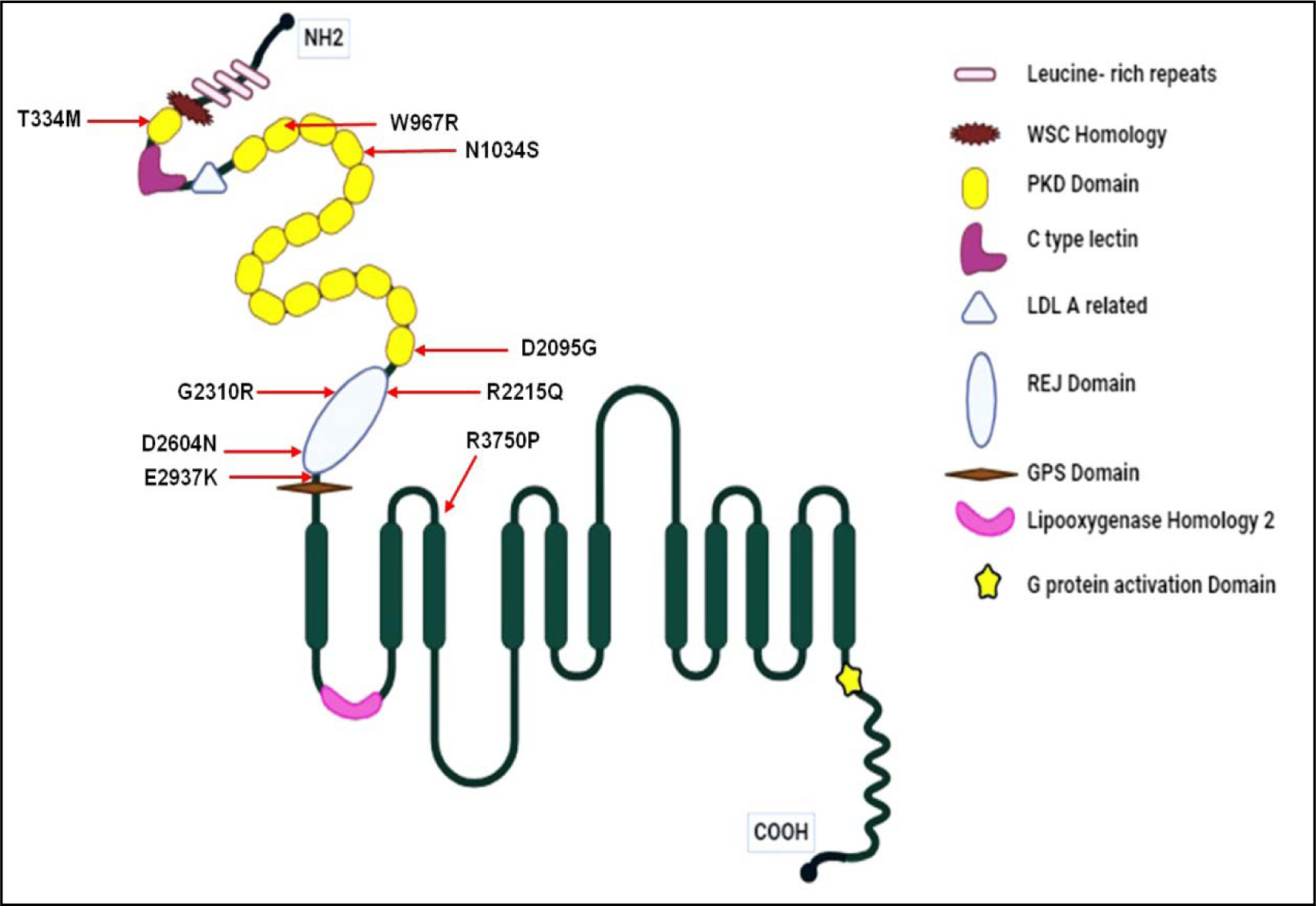
Structural Representation of *PKD1* Variants. The figure depicts the protein structure of polycystin-1 with marked missense variants analysed in this study. Each variant is marked within the protein structure, providing visual representation of their respective locations within the PC1 protein.

**Figure 2:**
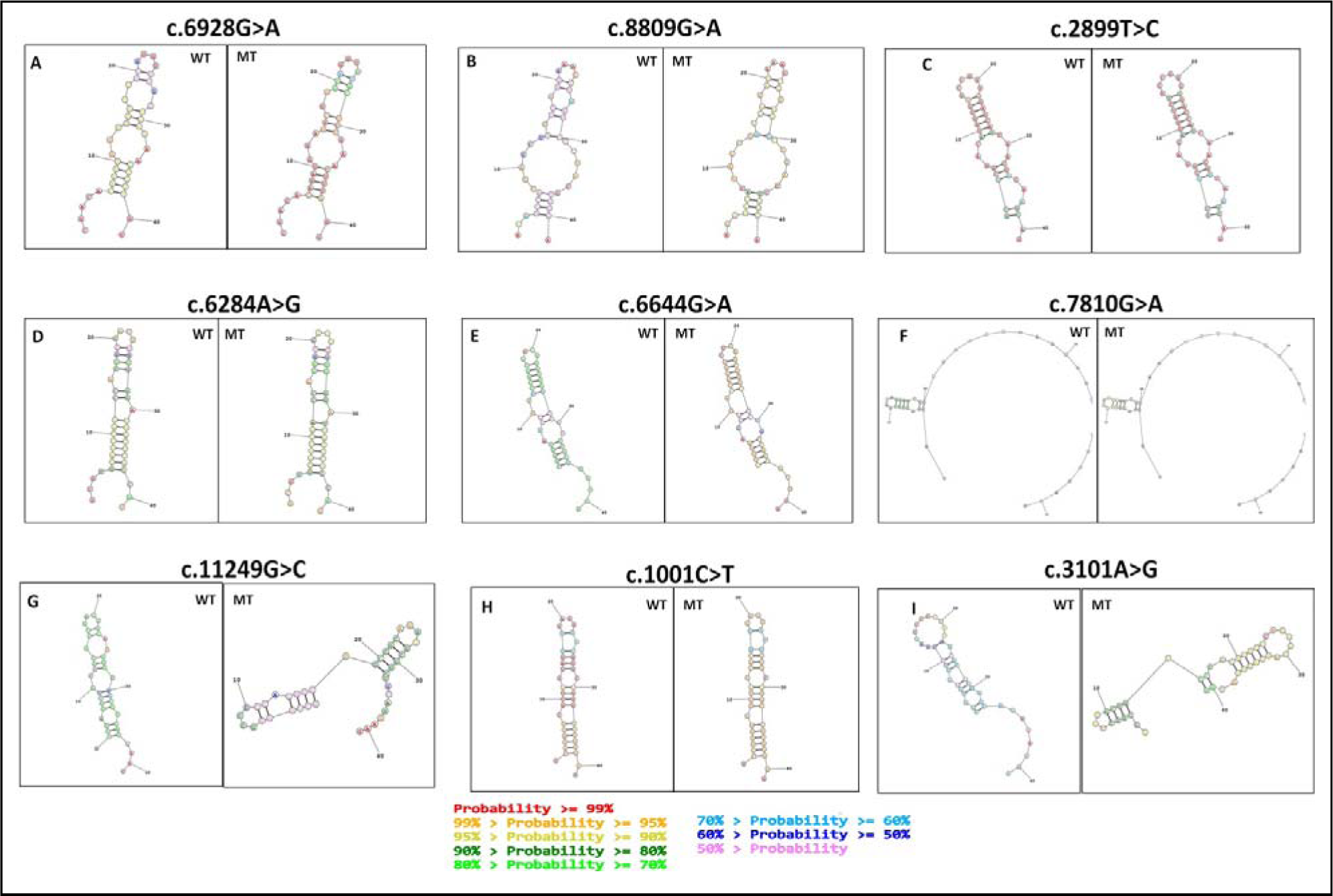
Secondary Structures of Original (WT) vs Altered (MT) RNA Snippet for *PKD1* Missense Variants. The cecondary structures of RNA snippets for each *PKD1* missense variants (from panels A to I) predicted using the RNAStructure Web server. The original (wild-type) RNA structure is depicted on the left, while the altered (mutant) RNA structure is shown on the right. The nucleotide change is at 21^st^ position in each structure Different colours of bases indicate the different base pairing probabilities.

### Assessment of Structural Impact using remuRNA

The *PKD1* missense variants were quantitatively assessed for their structural impact on RNA by analyzing the relative entropy H(wt:mu) (table 1). The greater the entropy value provided by remuRNA, the more is the structural impact of the variant (Miladi et al., 2020; Salari et al., 2013). This means that mutations with higher relative entropy values exhibit greater deviations in RNA structure compared to mutations with lower values. Variant c.6644G>A (p.R2215Q) exhibited the highest relative entropy value of 4.878, indicating substantial deviation in RNA structure induced by this mutation, followed by variants c.8809G>A (p.E2937K) and c.11249G>C (p.R3750P) with H(wt:mu) values of 4.644 and 4.642, respectively. Conversely, variants c.7810G>A (p.D2604N), c.6284A>G (p.D2095G), and c.2899T>C (p.W967R) showed relatively lower relative entropy values of 0.446, 0.148, and negligible 0.006, suggesting minimal structural impact on RNA. The remaining variants c.6928G>A (p.G2310R), c.3101A>G (p.N1034S), and c.1001C>T (p.T334M) exhibited moderate relative entropy values of 1.481, 2.912, and 0.192, respectively.

**Table 1:**
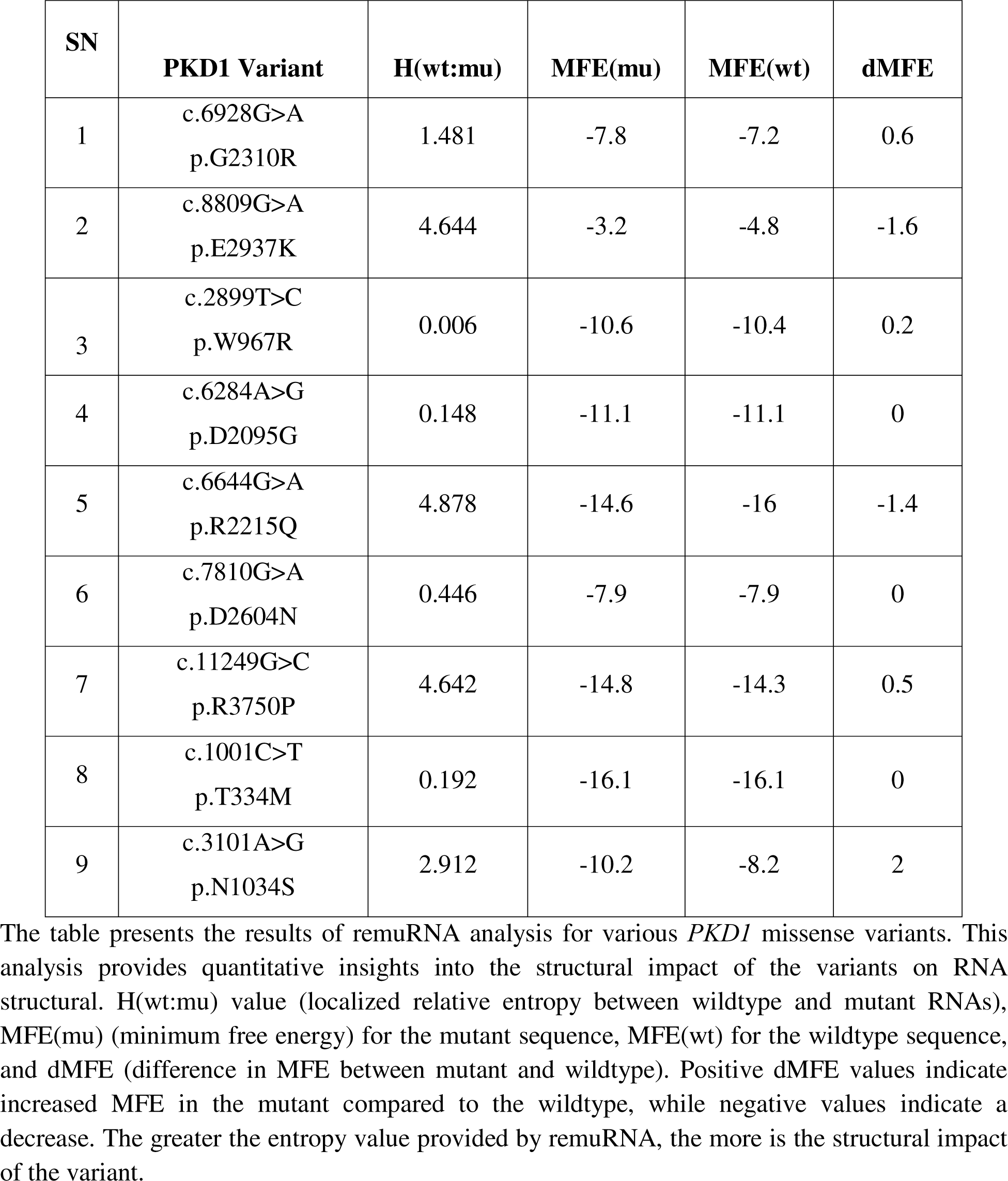
Quantitative Assessment of Structural Impact of *PKD1* Missense Variants on RNA.

| SN | PKD1 Variant | H(wt:mu) | MFE(mu) | MFE(wt) | dMFE |
| --- | --- | --- | --- | --- | --- |
| 1 | c.6928G>A<br>p.G2310R | 1.481 | -7.8 | -7.2 | 0.6 |
| 2 | c.8809G>A<br>p.E2937K | 4.644 | -3.2 | -4.8 | -1.6 |
| 3 | c.2899T>C<br>p.W967R | 0.006 | -10.6 | -10.4 | 0.2 |
| 4 | c.6284A>G<br>p.D2095G | 0.148 | -11.1 | -11.1 | 0 |
| 5 | c.6644G>A<br>p.R2215Q | 4.878 | -14.6 | -16 | -1.4 |
| 6 | c.7810G>A<br>p.D2604N | 0.446 | -7.9 | -7.9 | 0 |
| 7 | c.11249G>C<br>p.R3750P | 4.642 | -14.8 | -14.3 | 0.5 |
| 8 | c.1001C>T<br>p.T334M | 0.192 | -16.1 | -16.1 | 0 |
| 9 | c.3101A>G<br>p.N1034S | 2.912 | -10.2 | -8.2 | 2 |

The table presents the results of remuRNA analysis for various *PKD1* missense variants. This analysis provides quantitative insights into the structural impact of the variants on RNA structural. H(wt:mu) value (localized relative entropy between wildtype and mutant RNAs), MFE(mu) (minimum free energy) for the mutant sequence, MFE(wt) for the wildtype sequence, and dMFE (difference in MFE between mutant and wildtype). Positive dMFE values indicate increased MFE in the mutant compared to the wildtype, while negative values indicate a decrease. The greater the entropy value provided by remuRNA, the more is the structural impact of the variant.

The effect of the variants on RNA structure are illustrated using multiple approaches, including Circos plots (figure 3), base pairing probabilities dot plots (figure 4), differential base pairing probabilities dot plot (figure 5) and RNA accessibility profile analysis (figure 6). These MutaRNA findings are related to the effect of variant within the RNA snippet affecting the secondary structure and accessibility to its surrounding context.

**Figure 3:**
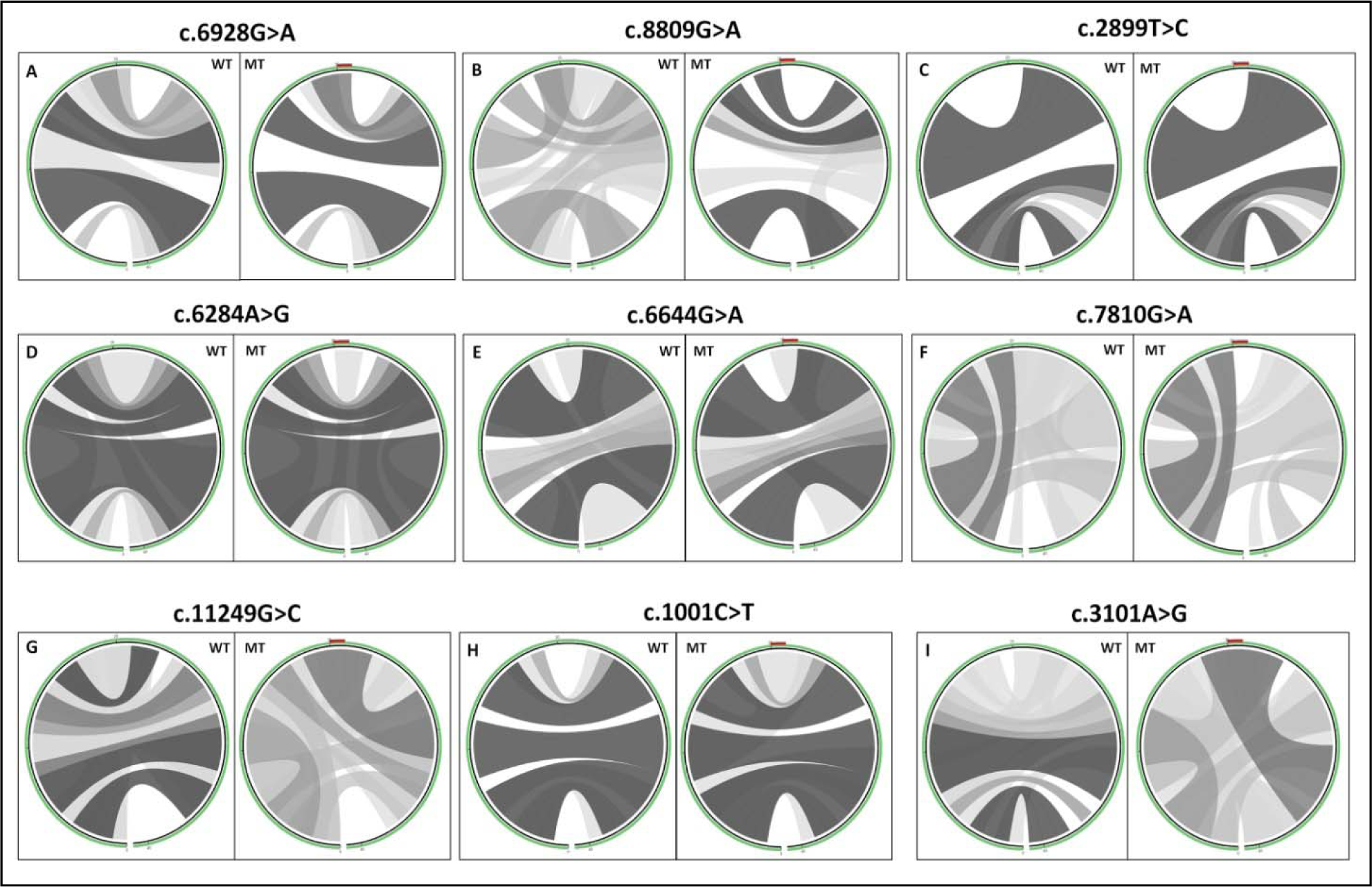
Circos Plots for Base Pair Probabilities of Original (WT) vs Altered (MT) RNA Snippets. The base pair probabilities are represented using circular plots (circos). The sequence starts from the 5’ end at the slight-left bottom and progresses clockwise until reaching the 3’ end. Within the circular plot, the analyzed mutation at position 21 is marked by a red color positioned at the top of the mutant circos. In each panel from A to I, the wild-type (WT) RNA structure is depicted on the left, while the mutant (MT) RNA structure is shown on the right. Darker hues of gray indicate higher probabilities of base pairing.

**Figure 4:**
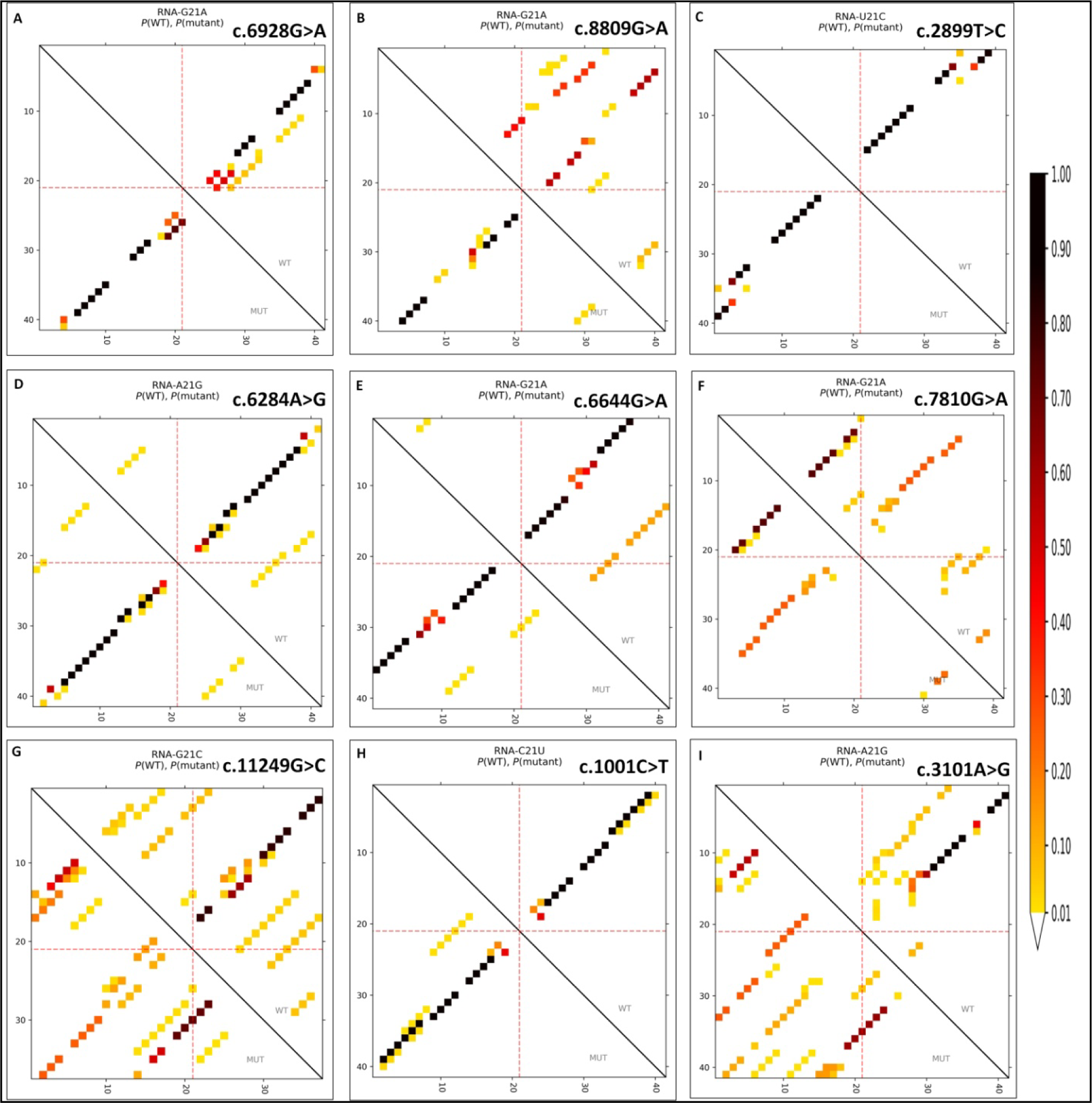
Base Pair Probabilities for Wildtype (WT) and Mutant (MT) *PKD1* Variants. The dot plot depicts the base pairing potential of both the WT and MT RNA variants. Darker dots indicate a higher probability of forming a base pair between the respective sequence positions. Base pair probabilities are derived from the Boltzmann distributed energies of all structures that can be formed by RNA, considering folding constraints such as maximal base pair span. The wildtype sequence is represented in the top-right portion of the matrix, while the mutant sequence is shown in the bottom-left. Each tick interval on the axes represents 10 nucleotides, with the evaluated variant at position 21 highlighted by red dotted lines on each axis. Panels A to I, represent each *PKD1* missense variants.

**Figure 5:**
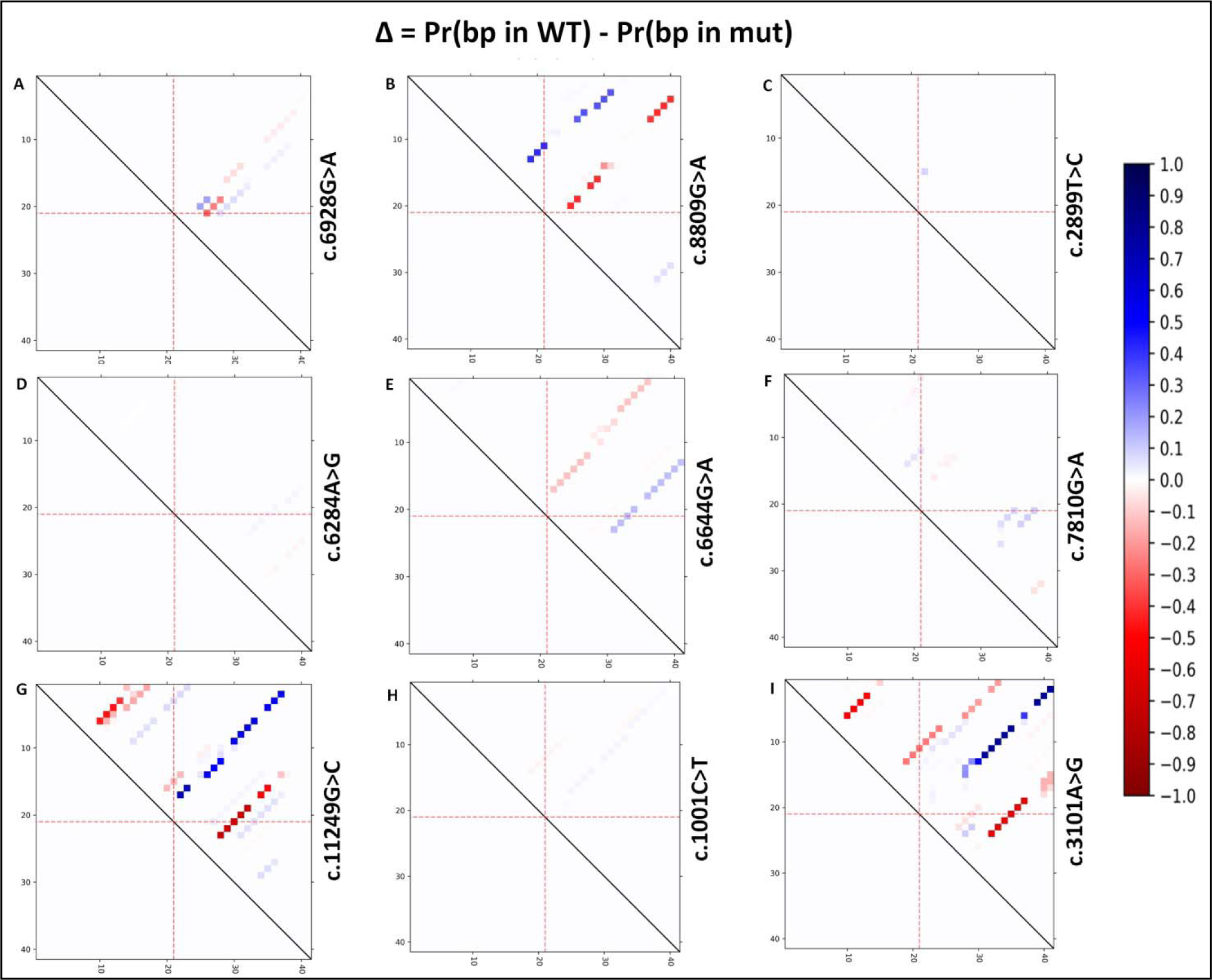
Differences in Base Pairing Probabilities Between Mutant and Wildtype RNA for PKD1 Missense Variants. The dot plot illustrates the difference in base pairing probabilities (Pr(bp)) between the mutant and wildtype RNA, calculated as Pr(bp in WT) - Pr(bp in mut) [Δ = Pr(bp in WT) - Pr(bp in mut)]. Weakened base pairs resulting from the mutation are represented in blue, while stronger ones are depicted in red. Each tick interval on the axes corresponds to 10 nucleotides. The mutated position is denoted by red dotted lines. Panels A-I represent individual *PKD1* missense variants.

**Figure 6:**
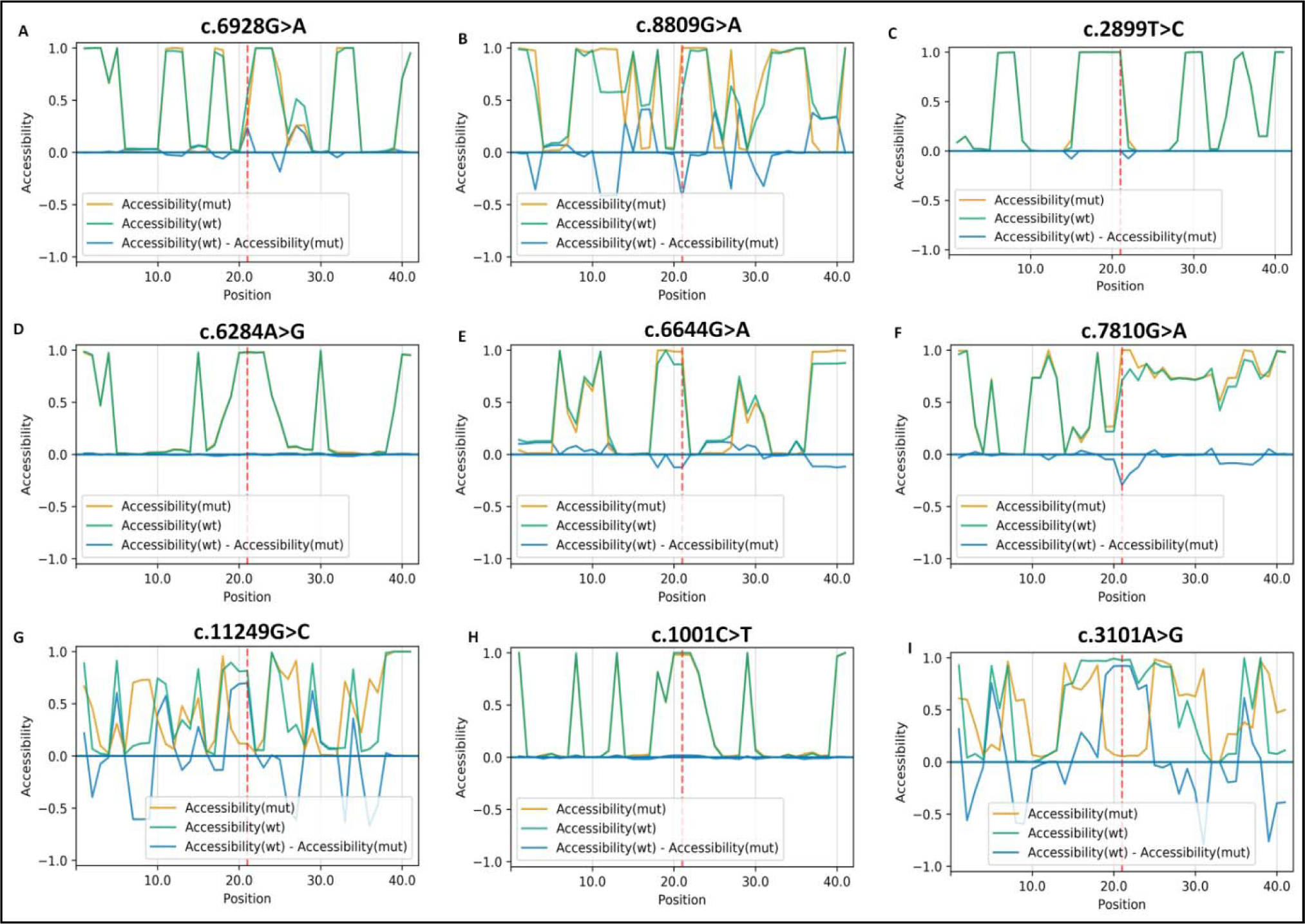
Comparison of Accessibility Profiles of Wild-Type (wt) and Mutant (mut) RNA Sequences for *PKD1* Missense Variants. The accessibility profiles of both wild-type and mutant sequences and their differences help evaluate how the mutation affects the RNA’s single-strandedness, which is strongly related to its interactions with other proteins or RNAs. Accessibility is evaluated in terms of local single-position unpaired probabilities, comparing the accessibility (i.e., probability of being unpaired) for each nucleotide position of the RNA sequences. The blue line represents the change in accessibility (WT-mut), with negative values indicating positions more likely to be unpaired in the mutant compared to the wild-type. This can be seen as the ‘negative drops’ in the blue differential accessibility profile. The red line indicates the mutated position for each *PKD1* missense variant (from panels A-I).

Circos Plots (Figure 3) were generated to visualize the interplay between different positions of the RNA sequence and their interactions. Variants were represented as arcs connecting the affected nucleotides, providing a view of the positional relationships within the RNA molecule highlighting potential disruptions caused by sequence change using different hues of grey colour with darker grey colour showing potentially stronger base pairing probabilities. Prominent deviations from wildtype patterns were observed in variants c.8809G>A (p.E2937K), c.11249G>C (p.R3750P), and c.3101A>G (p.N1034S) indicating significant structural impacts. Similarly, variants c.6928G>A (p.G2310R), c.6284A>G (p.D2095G), c.7810G>A (p.D2604N), exhibited less pronounced deviations, and c.2899T>C (p.W967R) and c.1001C>T (p.T334M) being the least, suggesting milder effects on RNA conformation. These base pair probabilities for wildtype (WT) and mutant (MT) sequences can also be visualized in heat maps dot matrices (Figure 4, 5). Consistent alterations in base pairing patterns were observed in variants indicating significant structural changes. This is also evident in the entropy chart and the secondary structures predictions using 2D structure and circus plots base pairing probabilities (table 1, figure 2 and 3).

The differences in base pairing probabilities between mutant and wildtype RNA (Δ = Pr(bp in WT) - Pr(bp in mut)) for *PKD1* missense variants are depicted as differential heat map dot matrices (Figure 5) to compare base pairing probabilities between mutant and wildtype RNA sequences with red indicating increased interaction likelihood and blue indicating weakened base pairs.

### RNA Accessibility

Comparison of accessibility profiles of wild-type (wt) and mutant (mut) RNA sequences for *PKD1* missense variants were compared to assess changes in single-strandedness and structure dynamics (Figure 6), which is strongly related to its interactions with other proteins or RNAs. The accessibility profile indicates the likelihood of nucleotides being unpaired, for each position within the RNA sequences. The blue line illustrates the alteration in accessibility (WT-mut), where negative values signify positions more likely to be unpaired in the mutant compared to the wildtype. The “negative drops” in the blue differential accessibility profile signifies the reduced accessibility in the folded wildtype compared to its mutant. The RNA accessibility profiles varied across different variants, with the prominent accessibility differences observed in variants c.8809G>A (p.E2937K), c.11249G>C (p.R3750P), and c.3101A>G (p.N1034S). Subsequently, c.6644G>A (p.R2215Q), c.6928G>A (p.G2310R), and c.7810G>A (p.D2604N) also exhibited effect on accessibility profile. Conversely, the accessibility was negligible in variants c.2899T>C (p.W967R), c.6284A>G (p.D2095G), and c.1001C>T (p.T334M). These findings suggest that certain variants, particularly c.8809G>A, c.11249G>C, and c.3101A>G (p.N1034S) may significantly impact RNA accessibility, potentially influencing interactions with other molecules or proteins. In contrast, variants such as c.6284A>G and c.1001C>T may have minimal effects on RNA accessibility, highlighting the variability in the structural consequences of the variants. The structural alterations in the predicted secondary structures are consistent with mutaRNA results.

### MD Simulation of protein structures

The RMSD, RMSF, SASA, Rg, and H-bonding plots for each variant with wild type are depicted in figure 7 to 11 respectively. The average values of each parameter are mentioned in table 2. For the c.8809G>A variant (p.E2937K), the mutant structure exhibited higher average RMSD (0.502 nm compared to 0.42 nm) and Rg (2.02 nm compared to 1.61 nm) values compared to the wild-type structure, indicating structural deviations and increased protein compactness. However, RMSF values remained relatively similar between the wild-type and mutant structures. The SASA and the number of hydrogen bonds were slightly higher in the mutant structure, suggesting changes in the protein’s surface area and hydrogen bonding interactions.

**Figure 7:**
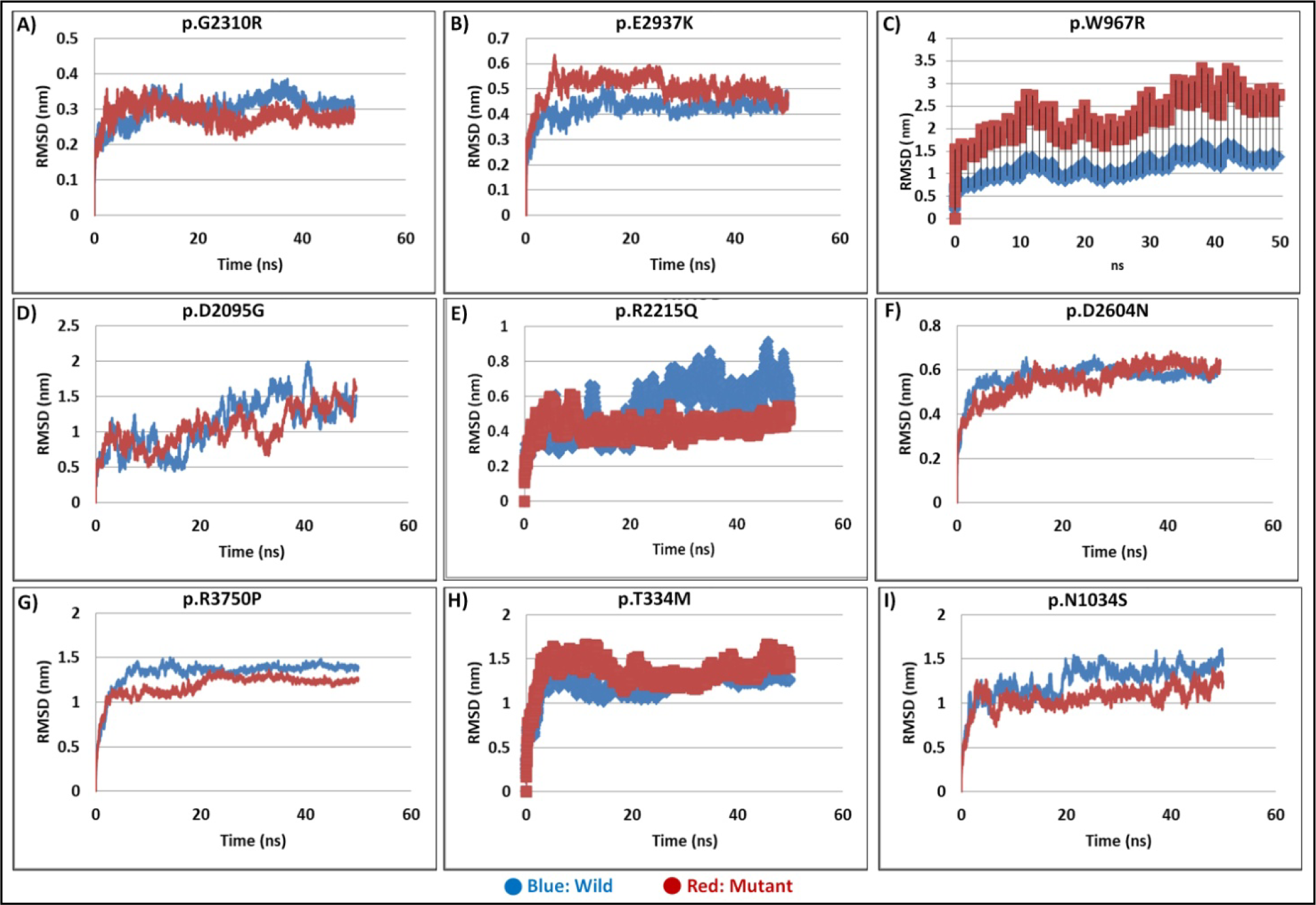
RMSD Analysis of the Variants: The RMSD plot depicts the average deviation of the backbone atoms of nine PKD1 variants vs wild-type from their initial structures over the 50ns time frame MD simulation. Each plot (A-I) represents the RMSD profile for a specific PKD1 variant (Red), with the wild-type (Blue). RMSD values were calculated relative to the initial conformation and plotted against simulation time (ns). The RMSD analysis predicts about the stability and structural changes of PKD1 variants compared to the wild-type protein.

**Table 2:**
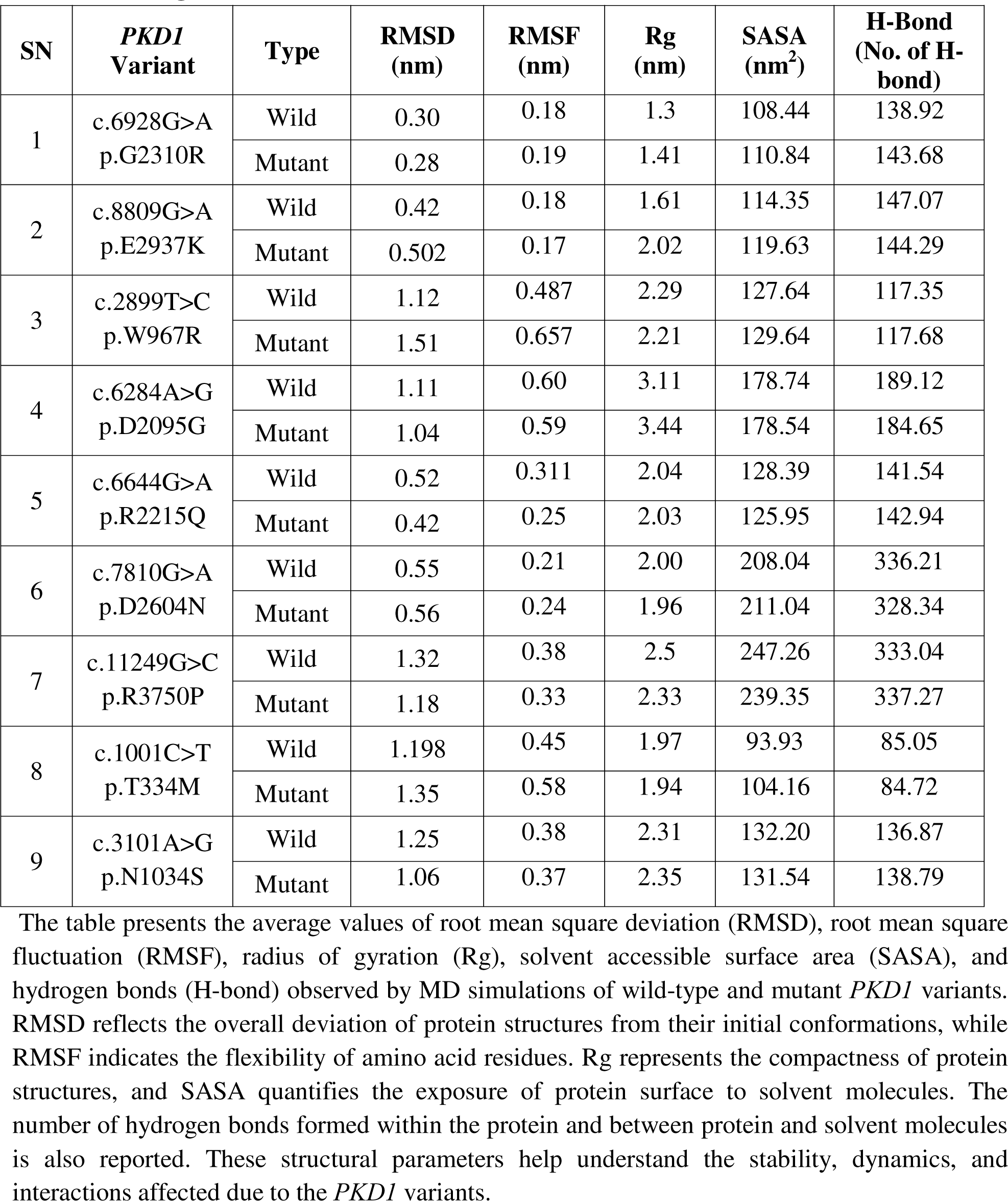
Average Values of Structural Parameters from MD Simulation.

For the c.6928G>A variant (p.G2310R), both wild-type and mutant structures exhibited differences in RMSD, with the mutant structure showing slightly lower RMSD (0.28 nm compared to 0.30 nm). RMSF values were comparable between the wild-type and mutant structures. However, the Rg value increased marginally in the mutant structure (1.41 nm compared to 1.3 nm in the wild type), indicating a slight expansion of the protein structure. Similarly, SASA and the number of hydrogen bonds also increased in the mutant structure, suggesting alterations in the protein’s surface characteristics and hydrogen bonding pattern.

For the c.2899T>C variant (p.W967R), considerable differences were observed between the wild-type and mutant structures. The mutant structure displayed higher RMSD (1.51 nm compared to 1.12 nm) and RMSF (0.657 nm compared to 0.487 nm) values, indicating increased structural flexibility and deviation from the native conformation. Rg values were comparable between the wild-type and mutant structures, while SASA and hydrogen bonding patterns showed minor variations.

For the c.6284A>G variant (p.D2095G), both wild-type and mutant structures exhibited similar RMSD and RMSF values. However, the mutant structure displayed a slightly higher Rg value (3.44 nm compared to 3.11 nm) and SASA, indicating a potential expansion of the protein structure and altered surface characteristics. The number of hydrogen bonds remained relatively stable between the wild-type and mutant structures.

For the c.6644G>A variant (p.R2215Q), the mutant structure showed slightly lower RMSD (0.42 nm compared to 0.52 nm) and RMSF values compared to the wild-type structure. Rg values were similar between the two structures, indicating comparable compactness. However, SASA was slightly reduced in the mutant structure, while the number of hydrogen bonds remained consistent.

For the c.7810G>A variant (p.D2604N), both wild-type and mutant structures exhibited similar RMSD and RMSF values. However, the mutant structure displayed a slightly higher Rg value (1.96 nm compared to 2.00 nm) and SASA, suggesting potential structural alterations and changes in surface accessibility. The number of hydrogen bonds was slightly lower in the mutant structure.

For the c.11249G>C variant (p.R3750P), the mutant structure exhibited slightly lower RMSD (1.18 nm compared to 1.32 nm) and RMSF values compared to the wild-type structure. Rg values were similar between the two structures, indicating comparable compactness. However, SASA was slightly reduced in the mutant structure, while the number of hydrogen bonds remained consistent.

For the c.1001C>T variant (p.T334M), the mutant structure displayed slightly higher RMSD (1.35 nm compared to 1.198 nm) and RMSF values compared to the wild-type structure. Rg values were similar between the two structures, indicating comparable compactness. However, SASA was slightly increased in the mutant structure, while the number of hydrogen bonds remained consistent.

For the c.3101A>G variant (p.N1034S), both wild-type and mutant structures exhibited similar RMSD and RMSF values. Rg values were also comparable between the two structures, indicating similar compactness. SASA and the number of hydrogen bonds remained relatively stable between the wild-type and mutant structures.

## Discussion

This study involves computational tools to investigate the effects on RNA structure and employed molecular dynamics simulation to explore the protein structure dynamics and functional implications of nine missense variants from the previously identified variants of PC1, the protein product encoded by the *PKD1*, recognized for its central role in ADPKD. The missense variants were individually identified in different ADPKD patients. ADPKD stands as the most prevalent hereditary renal disorder, characterized by the progressive formation of fluid-filled cysts within the kidneys, resulting in renal failure (Bergmann et al., 2018). PC1, a complex transmembrane protein, governs crucial cellular processes within renal tubules, considered vital for maintaining renal integrity (Paul & Vanden Heuvel, 2014; Weimbs, 2007). So far, the exact function of this protein largely remains elusive. The protein’s complexity arises from its large structure with multiple domains, each anticipated to contribute distinct functions. These include the leucine-rich repeat (LRR) domains, implicated in signal transduction and cell-matrix interactions, the C-type lectin domain facilitating protein-protein interactions, and the low-density lipoprotein-A (LDL-A) region, known for its cysteine-rich nature, however the presence of LDL-A in PC1 is still a debate. The 16 PKD repeats are considered to play essential roles in mediating cell-cell interactions and normal kidney development. The REJ domain possibly modulates ion transport, the PLAT domain facilitates protein-protein and protein-lipid interactions, and the 11 transmembrane domains likely act as ion transport channels. The cytoplasmic C-terminal tail regulates downstream signaling pathways by interacting with G protein subunits. Studying the dynamics of RNA structures *in vitro* is challenging and comprehending the role of PC1 in cellular physiology, particularly in the context of ADPKD, remains a formidable challenge, given its intricate nature as a large transmembrane protein with diverse domains (Wang et al., 2019; Weston et al., 2003). Through the analysis of various RNA structural parameters and MD simulation of protein regions may enable us to decipher the impact of mutations on RNA structure and interactions, PC1’s stability, dynamics, and interactions, offering helpful insights into the molecular foundations of ADPKD pathogenesis and paving the way for targeted therapeutic interventions.

The MD simulations conducted for various *PKD1* variants provided significant insights into the structural consequences of these mutations. As the protein size is large, motif and domain analysis through the motif scan web server aided in identifying variant locations and creating mutations within PKD1’s domain structure for simulation (DEVI et al., 2024). Analysis of RMSD, RMSF, Rg, SASA, and hydrogen bonding patterns revealed distinct differences between wild-type and mutant structures. For the c.6928G>A (p.G2310R), although the RMSD values were slightly lower in the mutant structure compared to the wild type, indicating a degree of stabilization, other parameters such as Rg, SASA, and hydrogen bonding showed modest increases (table 2; figure 7,9,10). This suggests that the mutation may induce subtle structural changes in the protein, potentially affecting its stability and surface properties. In contrast, the c.8809G>A (p.E2937K) exhibited notable deviations in RMSD and Rg values (table 2; figure 7, 10), indicating significant structural alterations in the mutant protein. The increased SASA and hydrogen bonding (figure 7, 11) observed in the mutant structure further emphasize the substantial impact of this mutation on the protein’s conformation and interactions. The c.2899T>C (p.W967R) showed the most noticeable differences, with substantially higher RMSD and RMSF values (table 2; figure 7 and 8) in the mutant structure, indicating increased flexibility and deviation from the native conformation. This mutation also resulted in a noteworthy increase in Rg and SASA, suggesting a more extended and exposed protein structure. For the c.6284A>G variant (p.D2095G), although the RMSD and RMSF values were comparable between wild-type and mutant structures, minor differences in Rg and SASA were observed, indicating alterations in the protein’s compactness and surface accessibility induced by the mutation. Similarly, the c.6644G>A variant (p.R2215Q) displayed slight differences in RMSD and RMSF values, with relatively stable Rg and SASA. These findings suggest that this mutation may also induce subtle changes in the protein structure without significantly affecting its overall compactness or surface properties. The c.7810G>A variant (p.D2604N) and c.11249G>C variant (p.R3750P) showed similar trends, with minor differences in RMSD and RMSF values but comparable Rg and SASA between wild-type and mutant structures suggesting that these mutations may have limited effects on the overall structure and surface properties of the protein. In contrast, the c.1001C>T variant (p.T334M) exhibited slightly higher RMSD and RMSF values in the mutant structure, indicating increased flexibility. However, other parameters such as Rg, SASA, and hydrogen bonding remained relatively stable, suggesting that this mutation may induce localized structural changes without significantly altering the overall protein conformation. For the c.3101A>G variant (p.N1034S), minimal differences were observed in RMSD, RMSF, Rg, SASA, and hydrogen bonding between wild-type and mutant structures, indicating that this mutation may have limited impact on the protein’s structure and dynamics. Thus, MD simulations provided valuable clues of the structural consequences of PKD1 mutations, highlighting their diverse effects on protein stability, flexibility, and surface properties.

**Figure 8:**
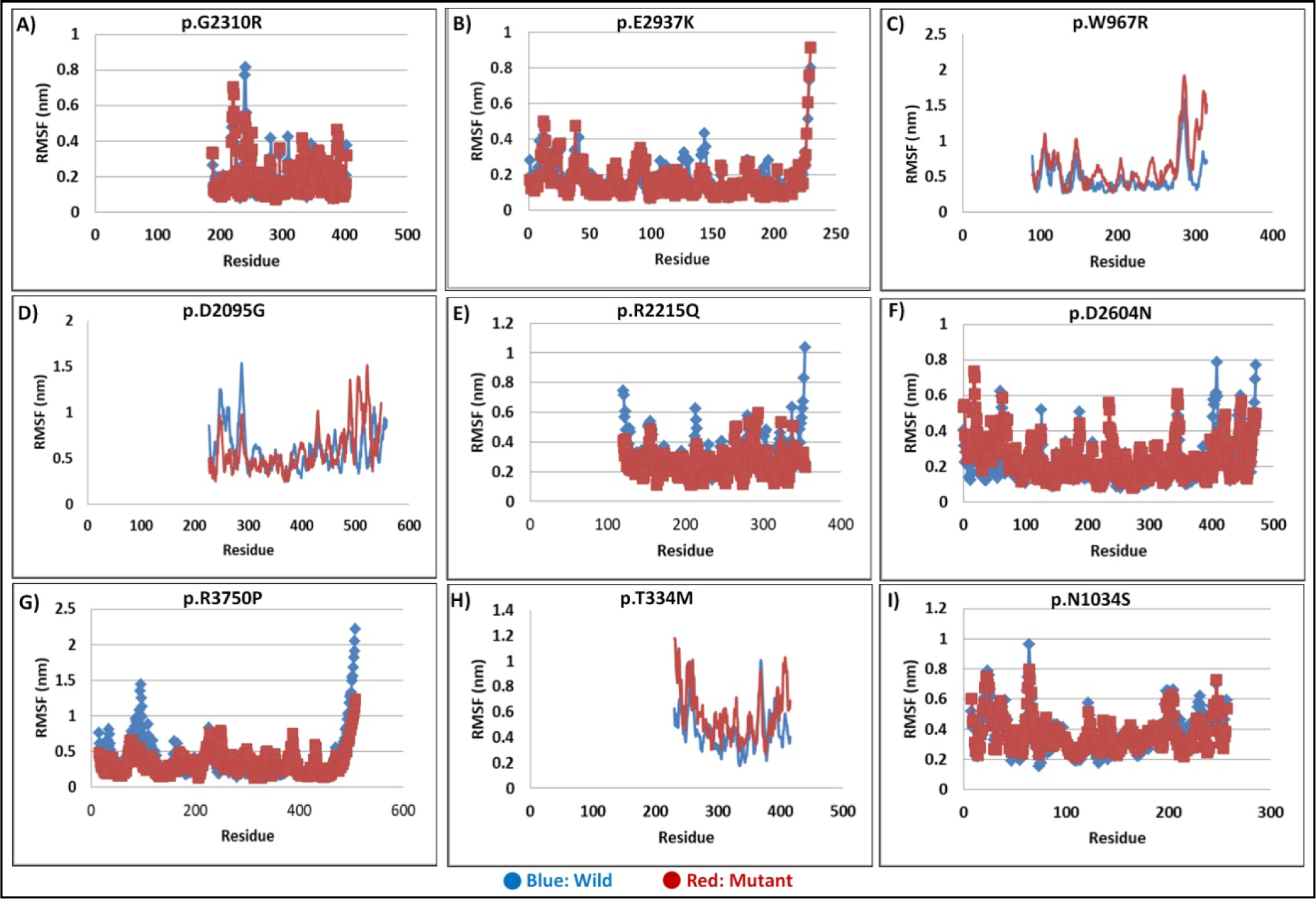
RMSF Analysis of PKD1 Variants: The RMSF plot illustrates the fluctuation of amino acid residues within the backbone of wild-type and mutant variants during 50ns time frame MD simulation. Each plat (A-I) represents the RMSF profile for a specific variant (Red), with the wild-type (Blue). The RMSF analysis provides insights into the individual residue dynamic flexibility and local structural variations of PKD1 variants compared to the wild-type protein.

**Figure 9:**
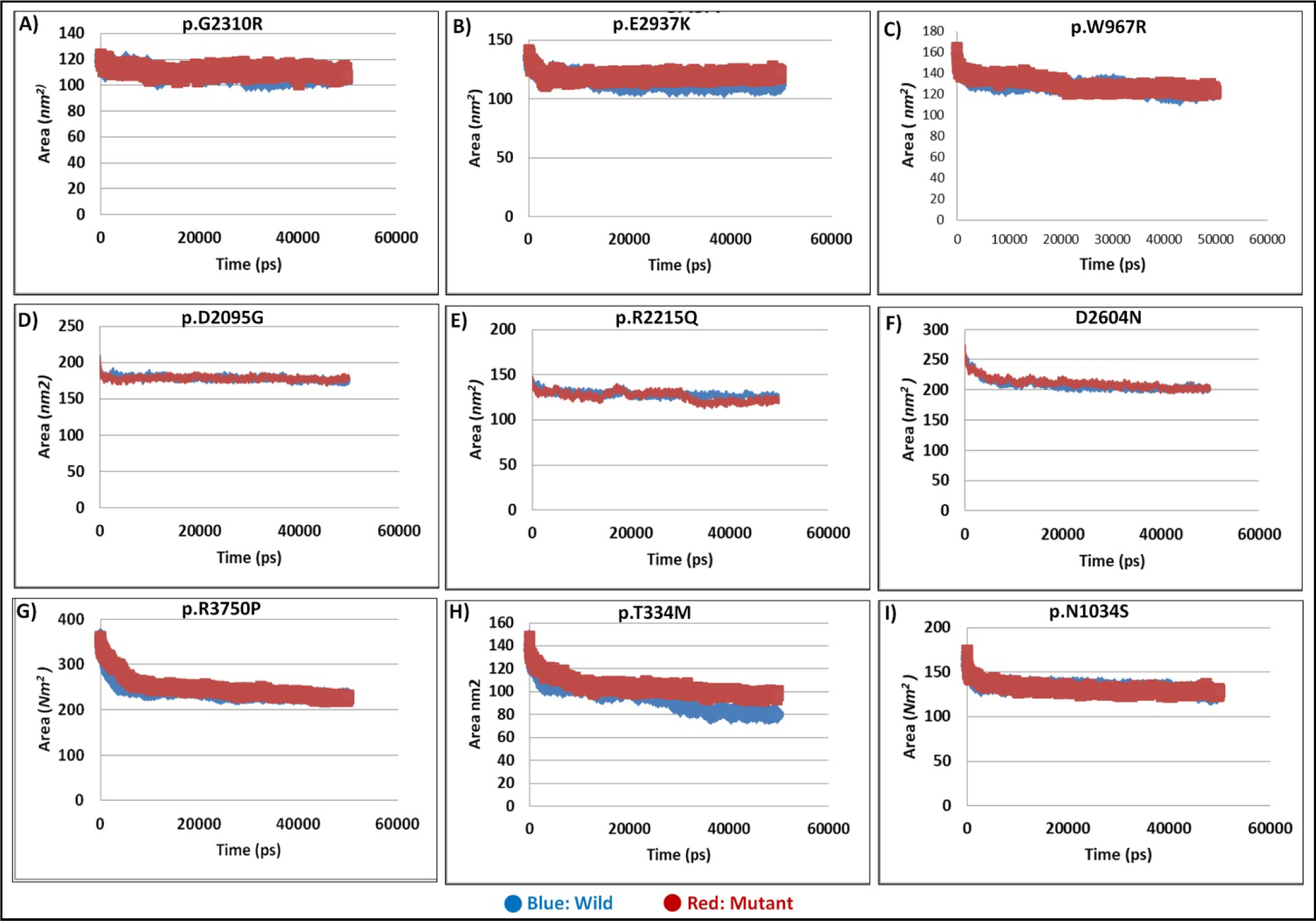
SASA Analysis of PKD1 Variants. The SASA plot displays the solvent accessible surface area of wild-type and mutant PKD1 variants over the 50 ns time frame. Each plot (A-I) represents the SASA profile for a specific PKD1 variant, with the wild-type depicted in a distinct color (Blue: wild, Red: mutant). The SASA analysis provides insights into the exposure of protein surfaces to solvent molecules, reflecting changes in protein interactions.

**Figure 10:**
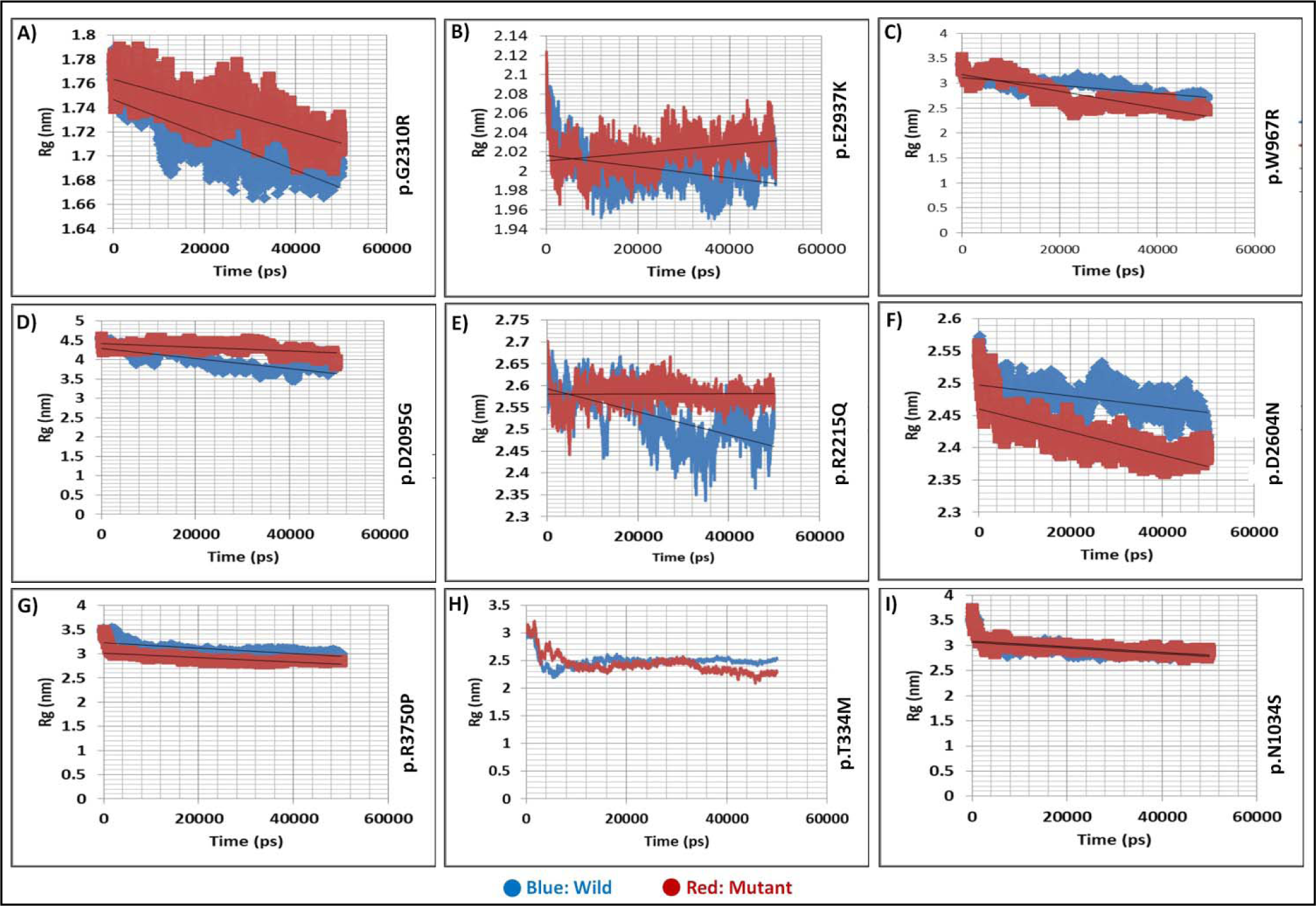
Radius of Gyration (Rg) Analysis of PKD1 Variants. The Rg plot illustrates the radius of gyration of wild-type and nine mutant PKD1 variants over 50ns time frame. Each plot (A-I) represents the Rg profile for a specific PKD1 variant (Red), with corresponding wild-type (Blue). Analysis of Rg profiles provides insights into the overall compactness of PKD1 variant protein compared to the wild-type protein.

**Figure 11:**
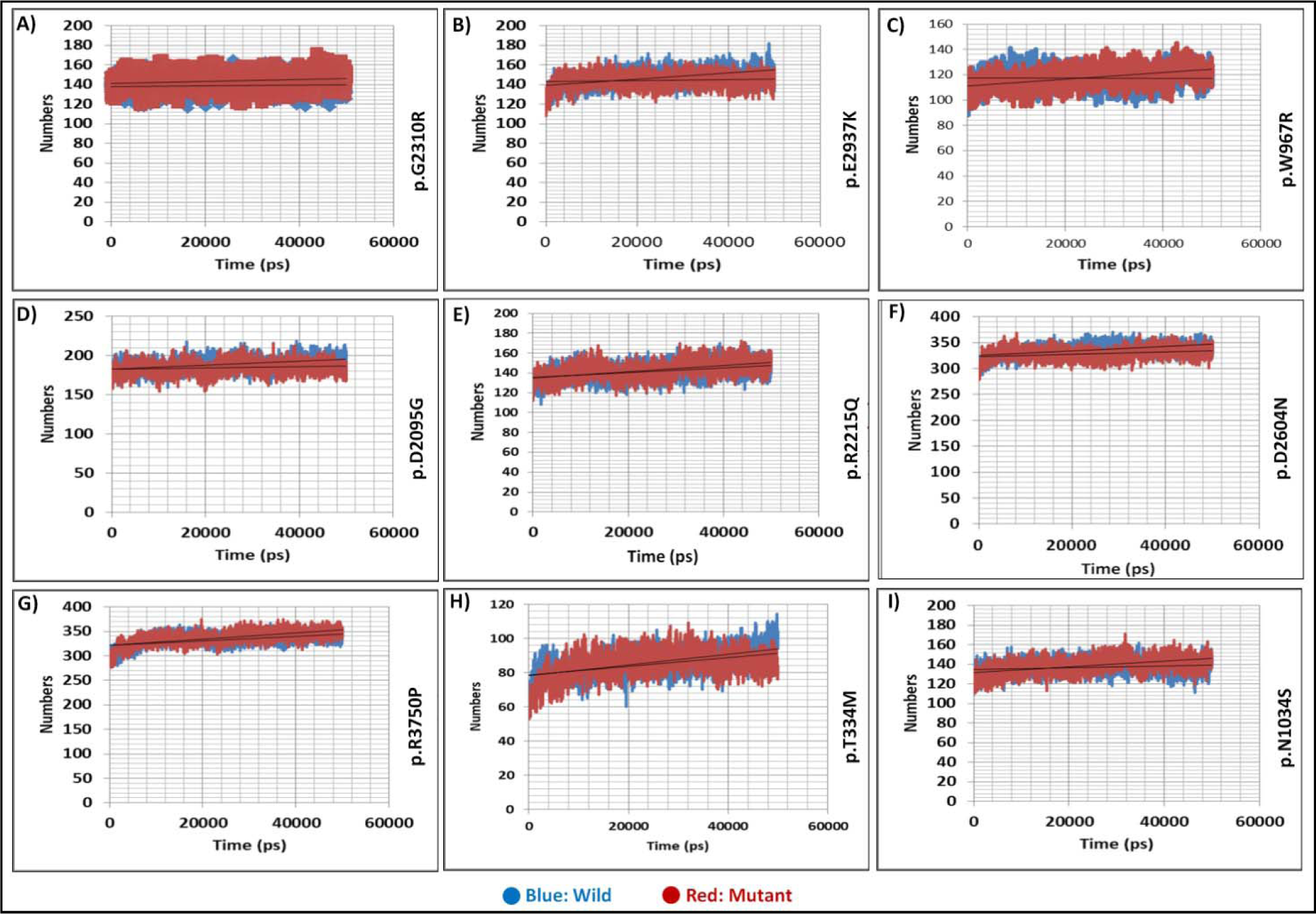
Hydrogen Bonding Analysis of PKD1 Variants. The H-bonding plot illustrates the number of hydrogen bonds formed within the protein and between protein and solvent molecules for wild-type and mutant PKD1 over 50ns time frame. Each plot (A-I) represents the H-bonding profile for a specific PKD1 variant (Red), with the corresponding wild-type (Blue). Analysis of H-bonding dynamics provides insights into the stability and interactions of PKD1 variants compared to the wild-type protein.

The studied missense variants in the *PKD1* are located in crucial domains involved in various aspects of PC1 function. The p.G2310R, p.R2215Q, p.D2604N, p.E2937K variant occurs within or near the REJ domain (figure 1), implicated in ion transport regulation, potentially altering its ability to modulate ion flux across cell membranes. The p.T334M, p.W967R, p.N1034S and p.D2095G variant in the 1^st^, 3^rd^, 5^th^ and 16^th^ PKD domain may impair PC1’s ability to interact with other proteins or structural elements. Similarly, the p.R3750P variant in the PKD cation channel domain may alter ion channel activity, affecting cellular ion homeostasis.

Talking about the RNA level, changes in secondary structure suggesting prominent or milder effects on RNA conformation highlight the heterogeneous nature of structural alterations induced by missense variants in *PKD1*, emphasizing the importance of considering the specific molecular consequences of each mutation in disease pathogenesis. The study comprehensively analyzed the structural impact of these variants on RNA secondary structure using a combination of computational tools and predictive models. RNA secondary structures were initially predicted using the RNAStructure web server, providing insights into the potential alterations induced by these variants. The analysis revealed prominent deviations from the wildtype RNA secondary structure in some variants which are c.6928G>A (p.G2310R), c.8809G>A (p.E2937K), c.6284A>G (p.D2095G), c.6644G>A (p.R2215Q), c.11249G>C (p.R3750P) and also c.3101A>G (p.N1034S). These variants exhibited significant impacts on RNA folding and stability, suggesting potential functional consequences at the molecular level. Further assessment of structural impact was analysed using the relative entropy between wildtype and mutant RNAs to quantify the extent of structural changes induced by each mutation (table 1). In descending order of relative entropy H(wt:mu) values, the most to the least impactful variants are as follows: c.6644G>A (p.R2215Q) with an H(wt:mu) of 4.878, c.8809G>A (p.E2937K) with 4.644, c.11249G>C (p.R3750P) with 4.642, c.3101A>G (p.N1034S) with 2.912, c.6928G>A (p.G2310R) with 1.481, c.7810G>A (p.D2604N) with 0.446, c.1001C>T (p.T334M) with 0.192, c.6284A>G (p.D2095G) with 0.148, c.2899T>C (p.W967R) with 0.006. These results indicate that variants such as c.6644G>A, c.11249G>C, and c.8809G>A have the most substantial impact on RNA structure, followed by c.3101A>G and c.6928G>A. Variants c.2899T>C (p.W967R) c.7810G>A, c.1001C>T, and c.6284A>G exhibit minimal effects on RNA conformation. These observed structural alterations were further corroborated by multiple approaches, including Circos plots, base pair probabilities dot plots, differential base pairing probabilities dot plots (figures 3, 4, 5). Similar trend was also observed in the RNA accessibility profile analysis with prominent differences observed in variants such as c.8809G>A (p.E2937K), c.11249G>C (p.R3750P), and c.3101A>G (p.N1034S). Variants like c.6644G>A (p.R2215Q), c.6928G>A (p.G2310R), and c.7810G>A (p.D2604N) also showed notable effects on the accessibility profile. In contrast, variants c.2899T>C (p.W967R), c.6284A>G (p.D2095G) and c.1001C>T (p.T334M) exhibited almost negligible changes in RNA accessibility. These findings suggest that certain variants, particularly in our study c.8809G>A, c.11249G>C and c.3101A>G (p.N1034S) may have a significant impact on RNA accessibility, potentially affecting interactions with other RNA or proteins (figure 6). The accessibility profile is evaluated in terms of probability of being unpaired for each nucleotide position of the RNA sequences. The accessibility profiles of both wild-type and mutant sequences and their differences help evaluate how the mutation affects the RNA’s interactions with other proteins or RNAs. These findings suggest the diversity in the structural consequences of the studied *PKD1* missense variants and the variants showing substantial structural alterations in RNA structure represent promising drug targets for precision medicine interventions and can be prioritized for future studies. The variants c.2899T>C (p.W967R), c.6284A>G (p.D2095G), c.7810G>A (p.D2604N), c.1001C>T (p.T334M) exhibited minimal alterations, suggesting milder effects on RNA level conformations, however, alterations were observed in MD simulations, particularly c.2899T>C (p.W967R) exhibited the most noticeable differences, telling that these variants may have effect on protein structure dynamics emphasizing the importance of considering both RNA and protein levels when assessing the functional consequences of genetic variants. Also, it is essential to note that even subtle structural changes induced by mutations can potentially alter RNA dynamics and function.

From this study we can infer that there is indeed a possibility that different missense variations can intricately influence RNA stability, structure, splicing patterns, translational efficiency, and protein structure dynamics differently and contribute to differences in disease presentation and progression, as observed in ADPKD. The study also highlights the heterogeneous nature of structural alterations induced by *PKD1* missense variants, drawing attention to the importance of considering the specific molecular consequences of each mutation in disease pathogenesis. We could deduce that the variant may impact RNA either alone or in conjunction with other genetic variants such as synonymous or frameshift variants. The synonymous variants, although not altering the amino acid sequence, can influence mRNA stability, ribosome binding, and translation kinetics, thereby impacting protein expression levels (Diederichs et al., 2016; Ganser et al., 2019). MutaRNA could come up as a useful tool to have a grasp on the structural consequences of synonymous variants on RNA (Miladi et al., 2020). These variants may disrupt RNA folding, alter splicing regulatory elements, impact translational kinetics, and act as disease modifiers by exacerbating the effects of other mutations, leading to a more severe disease phenotype (Diederichs et al., 2016). Moreover, the complex interplay between genetics, environmental factors, and epigenetic modifications can further modulate the overall impact of missense mutations on disease phenotypes. Understanding the multifaceted effects of missense variations on RNA is also essential for elucidating the molecular mechanisms underlying ADPKD pathogenesis. Insights from MD simulation provides additional grasp of how these missense variants could affect PC1 structure, stability, and interactions, aiding in understanding their functional consequences. While *in silico* tools cannot replace experiments conducted *in-vitro* or in model organisms, they can help sift through many variants to find a priority e.g., identifying regions of RNA or protein that undergo significant structural changes due to point mutations and guide experimental investigations. Simulation studies that examine RNA or protein structures in conditions resembling biological environments will improve our grasp of these structures within original physiological conditions.

By computationally examining the consequences of the missense variants at the RNA and protein level, we hope to deepen our understanding of molecular dynamics underlying ADPKD and identify new avenues for future therapeutic strategies targeting specific mutations. However, this study has several limitations as the scope of our investigation was limited to short snippets of RNA and specific regions/domains of the PC1 protein containing the identified variants. Though MD simulation offers valuable insights into protein dynamics and structure-function relationships, it inherently relies on computational modeling and predictions that are influenced by various factors, including the accuracy of force fields, simulation parameters, and initial protein structures, thus may not perfectly replicate biological reality. Hence our findings may not fully capture the comprehensive functional consequences of these mutations across the entirety of the protein; however it certainly gives initial close by insights in the dynamics affected by the point mutations. The experimental validation of our computational findings through biochemical and biophysical assays would fully weigh up the observations and establish the physiological relevance of the identified variants in the context of ADPKD.

## Conclusion

Through the systematic integration of computational methodologies, encompassing structural predictions, and MD simulations, this study analyzed *PKD1* missense variants structural and functional consequences at RNA and protein level. The analysis of parameters such as RMSD, RMSF, radius of gyration, SASA, and hydrogen bonding elucidated the effects of these variants on PC1 protein dynamics, stability, and interactions. The findings suggest that these variants may disrupt crucial domains such as the REJ domain, PKD domains, and cation channel domain, potentially compromising protein function. Variants including c.8809G>A (p.E2937K), c.11249G>C (p.R3750P), c.3101A>G (p.N1034S), c.6928G>A (p.G2310R), c.6644G>A (p.R2215Q) exhibited substantial alterations in RNA structures along with protein dynamics, suggesting prioritization for further functional implications as well as their potential as promising drug targets. We also observed that while some variants may not be influencing the RNA structure greatly but can affect the protein structure dynamics highlighting the importance of considering both RNA and protein levels while assessing their functional implications.

## Acknowledgements

We acknowledges the Senior Research Fellowship provided by the Indian Council of Medical Research (ICMR) to first author, India and Banaras Hindu University, Varanasi, India, for providing the internet and computer resources essential for conducting this study.

## References

Bellaousov, S., Reuter, J. S., Seetin, M. G., & Mathews, D. H. (2013). RNAstructure: web servers for RNA secondary structure prediction and analysis. Nucleic Acids Research, 41(W1), W471–W474. 10.1093/nar/gkt290

Bergmann, C., Guay-Woodford, L. M., Harris, P. C., Horie, S., Peters, D. J. M., & Torres, V. E. (2018). Polycystic kidney disease. Nature Reviews Disease Primers, 4(1), 50.

Bernhart, S. H., Mückstein, U., & Hofacker, I. L. (2011). RNA Accessibility in cubic time. Algorithms for Molecular Biology, 6, 1–7.

Butcher, S. E., & Pyle, A. M. (2011). The molecular interactions that stabilize RNA tertiary structure: RNA motifs, patterns, and networks. Accounts of Chemical Research, 44(12), 1302–1311.

Cowan, R., & Grosdidier, G. (2000). Visualization tools for monitoring and evaluation of distributed computing systems. Proc. of the International Conference on Computing in High Energy and Nuclear Physics, Padova, Italy.

Devi, C., Singh, S., Mohapatra, B., Kumar, A., Vikrant, S., Singh, R. G., Rai, P. K., & Das, P. (2024). A Whole Exome Sequencing Study of a small Indian Autosomal Dominant Polycystic Kidney Disease Patient Cohort. MedRxiv, 2023.04.20.23288719. 10.1101/2023.04.20.23288719

Diederichs, S., Bartsch, L., Berkmann, J. C., Fröse, K., Heitmann, J., Hoppe, C., Iggena, D., Jazmati, D., Karschnia, P., & Linsenmeier, M. (2016). The dark matter of the cancer genome: aberrations in regulatory elements, untranslated regions, splice sites, nonLJcoding RNA and synonymous mutations. EMBO Molecular Medicine, 8(5), 442–457.

Draper, D. E., Grilley, D., & Soto, A. M. (2005). Ions and RNA folding. Annu. Rev. Biophys. Biomol. Struct., 34, 221–243.

Ganser, L. R., Kelly, M. L., Herschlag, D., & Al-Hashimi, H. M. (2019). The roles of structural dynamics in the cellular functions of RNAs. Nature Reviews Molecular Cell Biology, 20(8), 474–489. 10.1038/s41580-019-0136-0

Halvorsen, M., Martin, J. S., Broadaway, S., & Laederach, A. (2010). Disease-associated mutations that alter the RNA structural ensemble. PLoS Genetics, 6(8), e1001074.

Holbrook, S. R. (2008). Structural Principles From Large RNAs. Annual Review of Biophysics, 37(1), 445–464. 10.1146/annurev.biophys.36.040306.132755

Hollingsworth, S. A., & Dror, R. O. (2018). Molecular dynamics simulation for all. Neuron, 99(6), 1129–1143.

Hopp, K., Cornec-Le Gall, E., Senum, S. R., Te Paske, I. B. A. W., Raj, S., Lavu, S., Baheti, S., Edwards, M. E., Madsen, C. D., & Heyer, C. M. (2020). Detection and characterization of mosaicism in autosomal dominant polycystic kidney disease. Kidney International, 97(2), 370–382.

Hunt, R. C., Simhadri, V. L., Iandoli, M., Sauna, Z. E., & Kimchi-Sarfaty, C. (2014). Exposing synonymous mutations. Trends in Genetics, 30(7), 308–321.

Miladi, M., Raden, M., Diederichs, S., & Backofen, R. (2020). MutaRNA: analysis and visualization of mutation-induced changes in RNA structure. Nucleic Acids Research, 48(W1), W287–W291.

Paul, B. M., & Vanden Heuvel, G. B. (2014). Kidney: polycystic kidney disease. Wiley Interdisciplinary Reviews: Developmental Biology, 3(6), 465–487.

Peintner, L., & Borner, C. (2017). Role of apoptosis in the development of autosomal dominant polycystic kidney disease (ADPKD). Cell and Tissue Research, 369, 27–39.

Raj, S., Singh, R. G., & Das, P. (2020). Mutational screening of PKD1 and PKD2 in Indian ADPKD patients identified 95 genetic variants. Mutation Research/Fundamental and Molecular Mechanisms of Mutagenesis, 821, 111718.

Ranjan, P., & Das, P. (2023). An inclusive study of deleterious missense PAX9 variants using user-friendly tools reveals structural, functional alterations, as well as potential therapeutic targets. International Journal of Biological Macromolecules, 233, 123375.

Salari, R., Kimchi-Sarfaty, C., Gottesman, M. M., & Przytycka, T. M. (2013). Sensitive measurement of single-nucleotide polymorphism-induced changes of RNA conformation: application to disease studies. Nucleic Acids Research, 41(1), 44–53. 10.1093/nar/gks1009

Salo-Ahen, O. M. H., Alanko, I., Bhadane, R., Bonvin, A. M. J. J., Honorato, R. V., Hossain, S., Juffer, A. H., Kabedev, A., Lahtela-Kakkonen, M., & Larsen, A. S. (2020). Molecular dynamics simulations in drug discovery and pharmaceutical development. Processes, 9(1), 71.

Sauna, Z. E., & Kimchi-Sarfaty, C. (2011). Understanding the contribution of synonymous mutations to human disease. Nature Reviews Genetics, 12(10), 683–691.

Schwede, T., Kopp, J., Guex, N., & Peitsch, M. C. (2003). SWISS-MODEL: an automated protein homology-modeling server. Nucleic Acids Research, 31(13), 3381–3385.

Sigrist, C. J. A., Cerutti, L., De Castro, E., Langendijk-Genevaux, P. S., Bulliard, V., Bairoch, A., & Hulo, N. (2010). PROSITE, a protein domain database for functional characterization and annotation. Nucleic Acids Research, 38(suppl_1), D161–D166.

Systèmes, D. (2016). Biovia, discovery studio modeling environment. Dassault Systèmes Biovia: San Diego, CA, USA.

Van Der Spoel, D., Lindahl, E., Hess, B., Groenhof, G., Mark, A. E., & Berendsen, H. J. C. (2005). GROMACS: fast, flexible, and free. Journal of Computational Chemistry, 26(16), 1701–1718.

Vander Meersche, Y., Cretin, G., Gheeraert, A., Gelly, J.-C., & Galochkina, T. (2024). ATLAS: protein flexibility description from atomistic molecular dynamics simulations. Nucleic Acids Research, 52(D1), D384–D392.

Wang, Z., Ng, C., Liu, X., Wang, Y., Li, B., Kashyap, P., Chaudhry, H. A., Castro, A., Kalontar, E. M., & Ilyayev, L. (2019). The ion channel function of polycystinLJ1 in the polycystinLJ1/polycystinLJ2 complex. EMBO Reports, 20(11), e48336.

Weimbs, T. (2007). Polycystic kidney disease and renal injury repair: common pathways, fluid flow, and the function of polycystin-1. American Journal of Physiology-Renal Physiology, 293(5), F1423–F1432.

Weston, B. S., Malhas, A. N., & Price, R. G. (2003). Structure–function relationships of the extracellular domain of the autosomal dominant polycystic kidney disease-associated protein, polycystin-1. FEBS Letters, 538(1–3), 8–13.

Xu, D., & Zhang, Y. (2011). Improving the physical realism and structural accuracy of protein models by a two-step atomic-level energy minimization. Biophysical Journal, 101(10), 2525–2534.

Yeung, K. C., Fryml, E., & Lanktree, M. B. (2024). How does ADPKD severity differ between family members? Kidney International Reports.

